# Redox regulation of K_v_7 channels through EF3 hand of calmodulin

**DOI:** 10.1101/2022.09.11.507486

**Authors:** Eider Nuñez, Frederick Jones, Arantza Muguruza-Montero, Janire Urrutia, Alejandra Aguado, Covadonga Malo, Ganeko Bernardo-Seisdedos, Carmen Domene, Oscar Millet, Nikita Gamper, Alvaro Villarroel

## Abstract

Neuronal K_V_7 channels, important regulators of cell excitability, are among the most sensitive proteins to reactive oxygen species. The S2S3 linker of the voltage sensor was reported as a site mediating redox modulation of the channels. Recent structural insights reveal potential interactions between this linker and the Ca^2+^-binding loop of the third EF-hand of calmodulin (CaM), which embraces an antiparallel fork formed by the C-terminal helices A and B. We found that precluding Ca^2+^ binding to the EF3 hand, but not to EF1, EF2 or EF4 hands, abolishes oxidation-induced enhancement of K_v_7.4 currents. Monitoring FRET between helices A and B tagged with fluorescent proteins, we observed that S2S3 peptides cause a reversal of the signal in the presence of Ca^2+^, but have no effect in the absence of this cation or if the peptide is oxidized. The capacity of loading EF3 with Ca^2+^ is essential for this reversal of the FRET signal, whereas the consequences of obliterating Ca^2+^ binding to EF1, EF2 or EF4 are negligible. Furthermore, we show that EF3 is necessary and sufficient to translate Ca^2+^ signals to reorient the AB fork. Our data is consistent with the proposal that oxidation of cysteine residues in the S2S3 loop relieves K_v_7 channels from a constitutive inhibition imposed by interactions between the EF3 hand of CaM which is necessary and sufficient for this signaling.

**Significance:** Oxidation-dependent enhancement of the K_V_7/M-channels plays a cytoprotective role in neurons. Here, we show that calmodulin (CaM), the main protein that conveys information from transient intracellular Ca^2+^ oscillations, plays a critical role in oxidative signal transduction. The prevailing view is that the main role of the EF-hands is to respond to Ca^2+^ and that the two EF-hands of CaM in each lobe act in coordination during signaling. We find that EF3 by itself is sufficient and necessary for the oxidative response of K_v_7 channel complex and for gating the Calcium Responsive Domain of K_v_7 channels. In addition, the direction of EF3-dependent signaling can be reversed by protein-protein interactions with solvent exposed regions outside the target binding groove between EF-hands.

## INTRODUCTION

The generation of abnormally high levels of reactive oxygen species (ROS) is linked to cellular dysfunction, including neuronal toxicity and neurodegeneration (3-5). In addition, ROS are important mediators of normal cellular functions in multiple intracellular signal transduction pathways (6-9). ROS generation induces oxidative modifications and augmentation of M-currents in neurons, which provides protective effects on oxidative stress-related neurodegeneration (10-12). K_V_7 channels, the substrate of the K_V_7-mediated M-current, are among the most sensitive proteins that respond to ROS production (3, 10, 13).

Superoxide anion radicals (O_2_•−), hydroxyl radicals (•OH), peroxynitrite (ONOO−), and hydrogen peroxide (H_2_O_2_) are the main ROS species produced in cells (14). These molecules display different reactivity, concentration and lifetime, and most probably play different roles in signal transduction and oxidative stress. Reversible oxidation of cysteine thiol side chains is one of the most recognized post-translational modifications produced by redox mechanisms in cells. Because of its relative stability and ability to cross the plasma membrane, H_2_O_2_ has been shown to be important in a variety of neurophysiological processes, including neurotransmission, ion channel function, and neuronal activity (15-18).

Augmentation of the M-current can be induced by an external H_2_O_2_ concentration as low as 5 μM (10), or even in the nM range (5). The M-current flows through channels formed of neuronal K_V_7 subunits (K_V_7.2-K_V_7.5, encoded by *KCNQ*2-5 genes). These tetrameric channels open at the subthreshold membrane potential and dampen cellular excitability (19, 20). K_V_7 channels have a core architecture similar to other voltage dependent potassium channels (1, 21-24): they have 6 helical transmembrane domains (S1-S6) with the voltage sensor formed by S1-S4, followed by a pore domain (S5-S6), which continues into a cytosolic C-terminal region. The C-terminus of K_V_7 channels contains five helical regions: helices A-D and TW helix between hA and hB. The latter region forms the Calcium Responsive Domain (CRD) with helices AB adopting an antiparallel fork disposition (25). Four C-helices from each subunit come together to form a stem perpendicular to the membrane. This stem continues with an unstructured linker that connects to helix D, which forms a tetrameric coiled-coil structure that confers subunit specificity during subunit assembly (25).

All K_V_7 channels require the association of calmodulin (CaM) to the CRD to be functional (26-28). Helices AB are embraced by CaM forming a compact structure just under the membrane that can move as a rigid body, with a region connecting S6 and helix A acting as a hinge (22-24). CaM is the main adaptor protein that confers Ca^2+^ sensitivity to an ample array of eukaryotic proteins, and is composed of two highly homologous lobes joined by a flexible linker. In solution, each lobe operates almost independently of the other and contains two similar Ca^2+^-binding EF-hands (29). This distinct signaling mediated by each CaM lobe was revealed early in *Paramecium* when it was discovered that mutations at the N-lobe affected a Ca^2+^ operated Na^+^ conductance, whereas mutations at the C-lobe affected a Ca^2+^ dependent K^+^ conductance (30).

A structure of the non-neuronal K_V_7.1 subunit trapped in a non-functional conformation with the voltage-sensor disengaged from the pore suggests that the EF3-hand of CaM may interact with the voltage-sensor (1, 23, 31) at a site essential for M-current redox modulation (5, 10). This 3D configuration has been assumed to confer a preferential use of EF3 during signaling on K_V_7 channels (31-34).

Here, we address the role of CaM on redox modulation of K_V_7 channels. We have monitored the role of each EF-hand in the redox-dependent current potentiation and in the gating process of the CRD, characterized by the opening of the AB fork (2). We find that the ability of EF3 to bind Ca^2+^ is critical for redox modulation of the CRD by S2S3 and for current potentiation, and this preferential signaling through EF3 is an intrinsic property of the CRD, not derived from the 3D arrangement observed in K_V_7 structures. Our data suggest that oxidation of K_V_7 channels enhances activity by disrupting the interaction between the S2S3 loop an EF3.

## RESULTS

### CaM plays a critical role in H_2_O_2_-mediated regulation of K_V_7.4 channels

We have previously shown that cysteine residues present in the unusual long intracellular linker between S2S3 transmembrane segments of K_V_7 channels are critical for H_2_O_2_-dependent potentiation (10). Recent studies suggest structural and functional interactions between this loop and calmodulin (CaM) (1, 31-34). To test a possible role of CaM in redox modulation, we used the perforated patch clamp method to measure K_V_7.4 activity in response to H_2_O_2_. Human *KCNQ4* cDNA was co-expressed in HEK293 cells with either CaM or CaM mutants that disable the Ca^2+^ binding ability of the N-lobe (CaM12), the C-lobe (CaM34), or both (CaM1234) (35, 36) (Fig 1). Bath-application of 150 µM H_2_O_2_ induced a clear augmentation of steady-state currents in the presence of CaM or CaM12 (Fig 1A-C). In contrast, the response was attenuated or precluded in the presence of CaM1234 or CaM34 (Fig 1A-C). Because structural and functional studies suggested a critical role of EF3 (1, 31-34), we tested the effect of CaM3 and CaM124. Whereas the H_2_O_2_ response in the presence of CaM124 (Fig 1A-C) was maintained, it was diminished with CaM3 (Fig 1D). Importantly, while the response to H_2_O_2_ was abolished in the presence of CaM3, another K_V_7 activator, retigabine, still produced strong activation of K_V_7.4 current under these conditions (Fig 1D). Retigabine activates K_V_7 channels by binding to a hydrophobic pocket between S4 and S5 domains, a site that does not overlap with CaM binding site (37). These results suggest that EF3 of CaM is necessary for augmentation of K_V_7 channels by H_2_O_2_ specifically.

**Figure 1.**
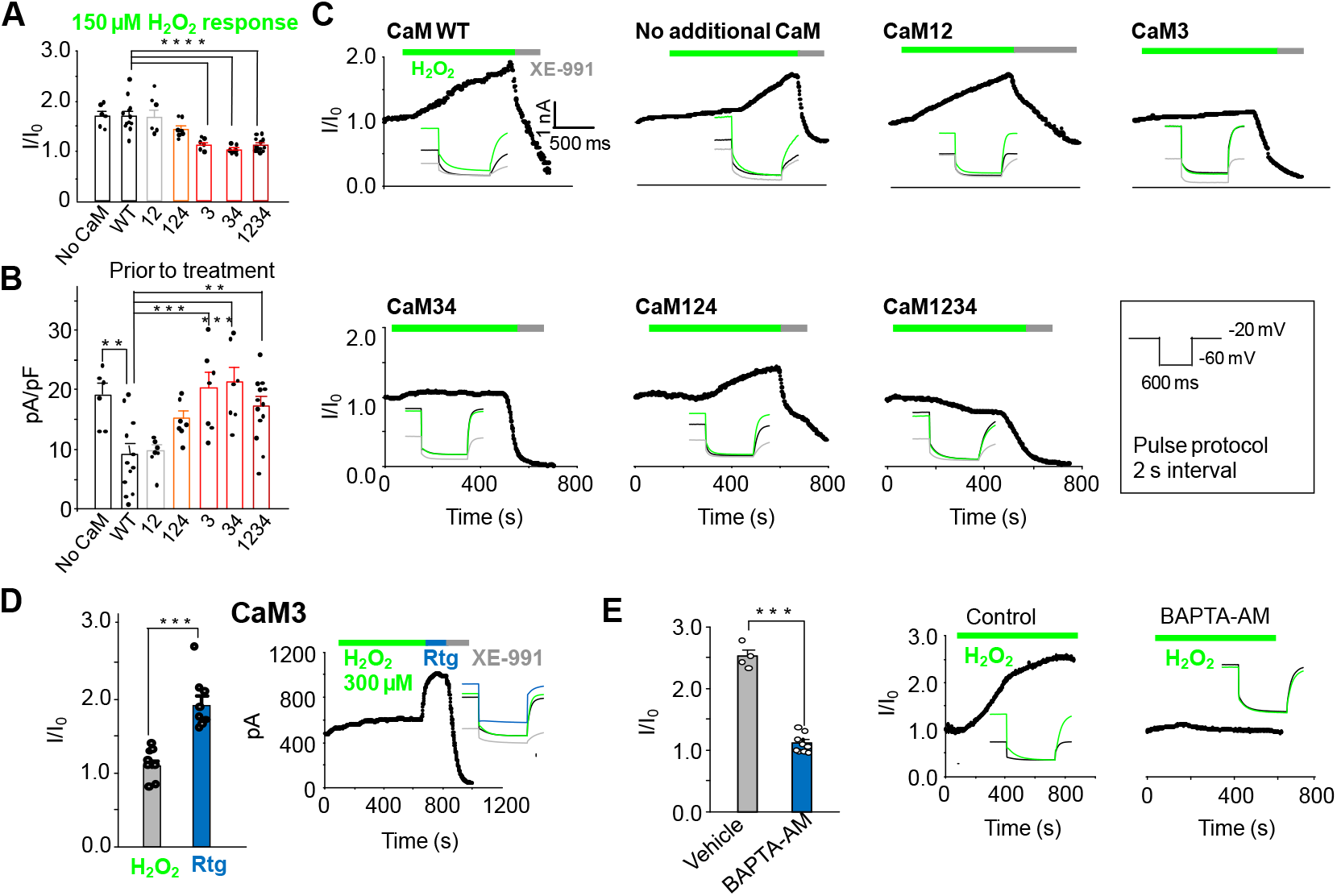
EF-3 hand Ca^2+^ binding capacity of CaM is required for H_2_O_2_ mediated potentiation of K_V_7.4. **A**. Response of K_V_7.4 transfected HEK293 cells to 150 µM H_2_O_2_ (I/I_0_) when co-transfected with wild-type CaM (CaMWT n = 12) or mutant CaMs lacking Ca^2+^ binding to one or more EF hands. The number in X-axis of pertains to the EF hand unable to bind Ca^2+^ (CaM12 n = 7, CaM124 n = 7, CaM3 n = 7, CaM34 n = 7 and CaM1234 n = 13) or with no additional CaM transfected (No CaM, n = 6). **B**. Current density (pA/pF) of K_V_7.4 transfected cells prior to treatment with H_2_O_2_. **C**. Example time courses of the effects of H_2_O_2_ (150 µM) and of the specific M-channel blocker, XE-991 (10 µM), on the outward steady-state current at -60 mV in HEK293 cells overexpressing K_V_7.4 and one of the CaM variants. The voltage protocol is shown in a black box. **Insets:** representative current traces from each condition (control: black trace, H_2_O_2_ green trace, XE-991: grey trace). **D**. Comparative response of cells transfected with K_V_7.4 and CaM3 to 300 µM H_2_O_2_ and 10 µM retigabine (n = 8). **Insets:** representative traces for control (black), H_2_O_2_ (green), retigabine (blue) and XE-991 (grey). **E**. Ca^2+^ dependence of H_2_O_2_ response in cells transfected with K_V_7.4 and CaMWT. Comparison of 300 µM H_2_O_2_ response in normal or low Ca^2+^ conditions induced by pre-incubation of cells in 10 µM BAPTA-AM for 30 minutes to chelate intracellular Ca^2+^. Control n = 4, BAPTA-AM n = 9. **Insets:** representative traces in control (black) and H_2_O_2_ (green). Data presented is mean ± SEM, statistical evaluation by independent measures ANOVA with Dunnett’s post hoc, **P<0.01, ***P<0.001, ****P<0.0001 (A&B). A paired (D) or unpaired (E) two-tailed T test ***P<0.001, ****P<0.0001.

EF-hand mutations used above mimic Ca^2+^-free (apo) state of the CaM, with CaM1234 being completely Ca^2+^-free, while other mutants are partially Ca^2+^-free. Since CaM1234 prevented the K_V_7.4 current augmentation by H_2_O_2_ (as did the other mutants containing EF3 mutation), we therefore tested if ‘sponging’ intracellular Ca^2+^ by pre-incubating the cells with BAPTA-AM also prevent the H_2_O_2_ effect on K_V_7.4. BAPTA-AM crosses the membrane, and release the strong Ca^2+^ chelator BAPTA intracellularly, thereby lowering resting free Ca^2+^ levels. The response to oxidation was indeed virtually abolished under these conditions.

As expected, the effect of H_2_O_2_ was absent after substituting the redox-sensitive triplet of cysteine residues at the positions 156, 157, 158 in the S2S3 linker of K_V_7.4 by alanine residues (Supplemental Fig 2). There was no difference whether WT CaM or CaM1234 was present, in either case the current produced by CCCAAA K_V_7.4 was only marginally affected by 150 µM H_2_O_2_ (Supplemental Fig 2). Interestingly, all CaM mutants containing EF3 mutations (CaM3, CaM34 and CaM1234) produced small negative shift in K_V_7.4 voltage dependence (Supplemental Fig 1), a finding consistent with a presumed removal of a tonic inhibitory effect of calcified EF3.

**Figure 2.**
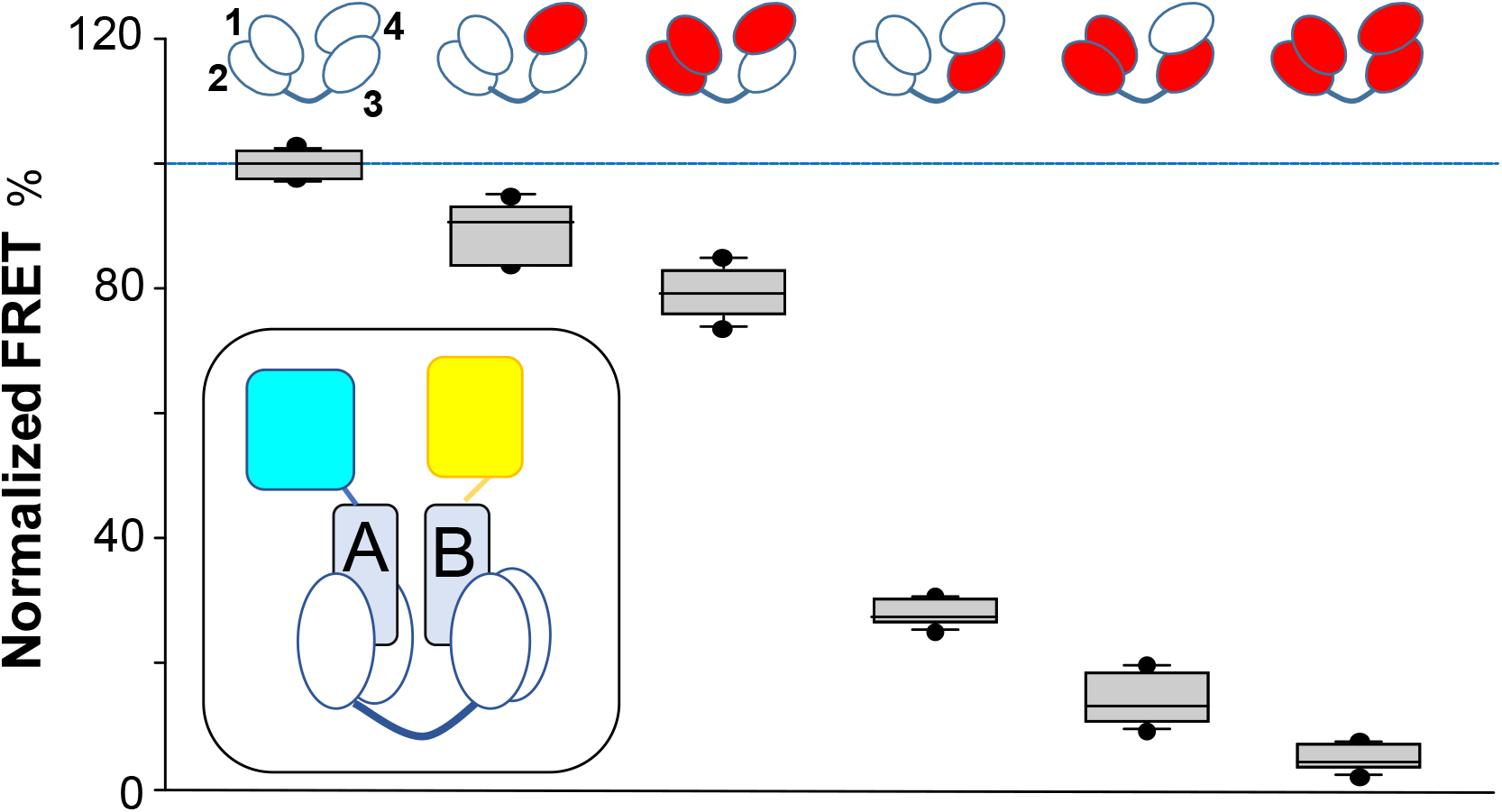
Influence of EF hand mutations abolishing Ca^2+^-binding to CaM on Ca^2+^-dependent interaction between CaM and the AB fork of K_V_7.2, as measured by FRET. Top. Cartoon representation of CaM mutants. The EF-hands carrying a mutation that preclude Ca^2+^ binding are colored in red. Bottom, box-plot of the relative FRET index change produced by Ca^2+^ for the AB fork in complex with the indicated mutated CaM. Note that in the complex with CaM3 and CaM123 the changes prompted by Ca^2+^ were almost obliterated, whereas in the complex with CaM124 the response was preserved. Each plot represents the average of 6 independent experiments. **Inset:** Cartoon representing the FRET sensor in complex with CaM (mTFP1-hA-hB-Venus/CaM).

Overall, these experiments suggested that EF3 of CaM and cysteine residues in the S2S3 of K_V_7.4 are necessary for current activation by H_2_O_2_. We hypothesize that binding of Ca^2+^ to EF3 partially inhibits K_V_7.4; preventing such binding or removing Ca^2+^ from this location disinhibits the channel. We further hypothesize that oxidative modification of S2S3 cysteine residues antagonizes the EF3/Ca^2+^ inhibition of K_V_7.4. To get further insights, we analyzed the behavior of the isolated CRD, without constrains imposed by other channel domains, the membrane or the complexity of potential intracellular signaling cascades evoked *in vivo*.

### Ca^2+^ binding to EF3 is critical for signaling within the Calcium Responsive Domain

Wild-type or mutant CaMs were co-expressed with the CRD from all five K_V_7 subunits in bacteria, the complexes were purified, and Ca^2+^ signaling was examined by monitoring the transfer of energy between the two fluorophores attached to the N- and C-termini of the AB fork (See inset in Fig. 2). Experiments with AB fork of K_V_7.2 are shown in Figure 2 and results for CRD from other K_V_7 subunits are shown in Supplemental Figure. 4. CaM remained firmly attached to the AB fork under our *in vitro* conditions (Supplemental Fig 3). FRET efficiency was reduced in a Ca^2+^ concentration-dependent manner as previously described (2). Mutations into EF1 and EF2 (CaM12) did not significantly alter Ca^2+^-dependent signaling (n = 6), whereas mutations at either EF-hands 3 or 4 (CaM3 or CaM4) reduced the magnitude of FRET changes (n = 6). The extent of the effect was significantly decreased in the complex with CaM3, with a minor effect in the complex with CaM4 (Fig 2 and Supplemental Fig 4).

**Figure 3.**
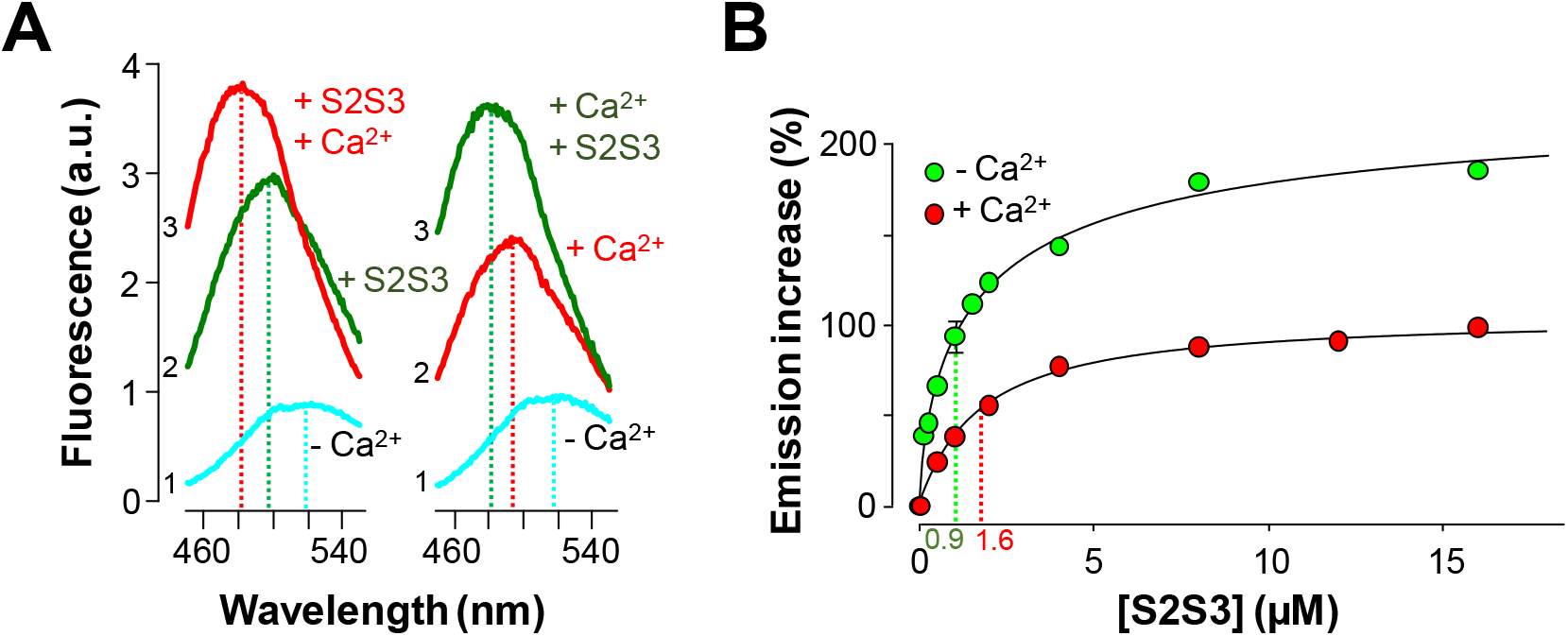
Effects of a 24 residues K_v_7. S2S3 peptide on fluorescence emission of dansylated calmodulin. **A**. Emission spectra of D-CaM (50 nM) in Ca^2+^-free conditions (cyan), and after subsequent sequential addition of the S2S3 peptide (16 µM, green), and Ca^2+^ (10 µM free concentration, red). The order of additions is indicated at the left of each trace. **B**. Dose-dependent relative fluorescent emission increase as a function of S2S3 peptide concentration, in the absence (green) and the presence of Ca^2+^ (10 µM, red). A Hill equation was fitted to the data (continuous line) with EC_50_ = 0.88 ± 0.12 and 1.63 ± 0.07 µM (n ≥ 6), in the absence and the presence of Ca^2+^, respectively. The K_V_7.1 S2S3 peptide sequence was Ac-RLWSAGCRSKYVGVWGRLRFARK-NH_2_. Similar results were obtained for K_V_7.1 through K_V_7.5 peptides (Supplemental Fig 5).

**Figure 4.**
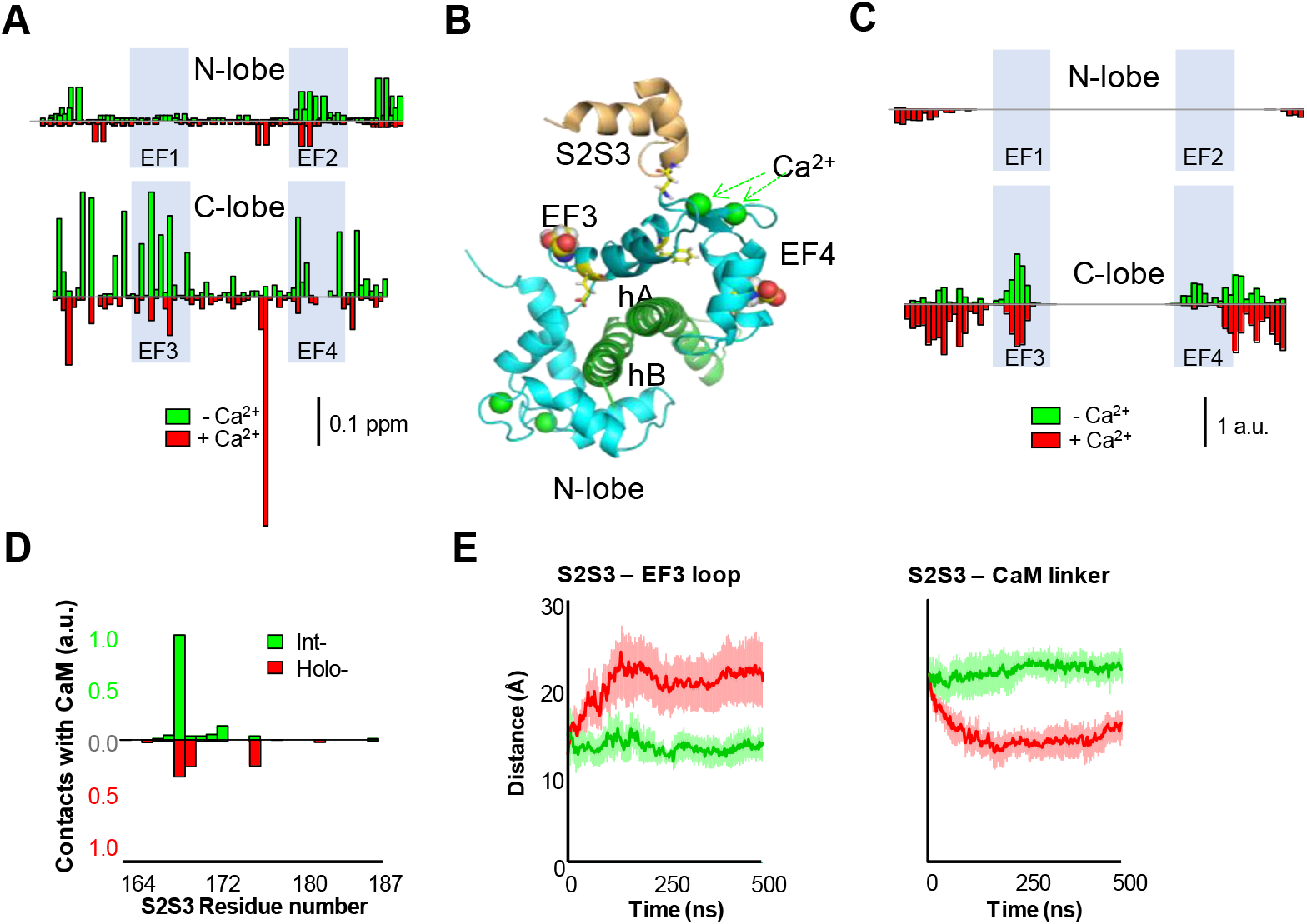
Interaction between the K_V_7 S2S3 peptide and CaM in complex with K_V_7.2 CDR. **A**. The CSP analysis shows that the magnitude of local residue environmental alterations detected by NMR are lager in the C-lobe, both in the presence and in the absence of Ca^2+^. **B**. Structural mapping of the main CSPs in the presence of Ca^2+^ over Ca^2+^-loaded K_V_7.2 CaM/CDR complex. Residues with displacement above 3 times the mean are shown as sticks, and the two residues with the larger displacements are represented as balls. The structure of the S2S3 loop was derived from the Cryo-EM PDB 5VMS (1) and placed according to structural alignment of the C-lobe of PDB 6FEH (2). **C**. Contact map derived from MD simulations of the S2S3/CaM complex. Normalized CaM contacts with the S2S3 peptide residues (10 Å cut-off) for int-(green) and holo-systems (red) (See Supplementary Fig 9). Vertical calibration bar is in arbitrary units (a.u.). **D**. S2S3 contact map with CaM residues (4 Å cut-off)(see Supplementary Fig 9). **E**. Distance as a function of time between the mass centers of the EF3 loop (residues D93-G98) and (i) the S2S3 loop (residues R164-L173) (left) or (ii) the linker connecting CaM lobes (residues R74-E84) (right). Bars indicate SEM (n = 6).

The role of EF3 was further examined combining Ca^2+^-binding canceling mutations in EF1, EF2 and EF4-hands. The AB/CaM124 complex, that is, with only EF3 able to bind Ca^2+^, presented a response to Ca^2+^ that was ∼80% that of the AB in complex with WT CaM (n = 6). In contrast, the response of the complex with CaM3 was reduced to ∼30% (n = 6). A similar strategy was followed to evaluate the role of the EF4 hand, testing complexes with CaM123 and CaM4. In the complex with CaM123, the response was almost abolished, whereas in the complex with CaM4 the response was about 90% of that of WT (n = 6) (Fig 2). Thus, EF3 plays a significant role in transmitting Ca^2+^ signals to the AB fork, and EF4 plays a secondary function.

### Peptides derived from the K_V_7 S2S3 loop interact with CaM

A subset of cryo-EM K_V_7.1 channel particles has revealed a likely interaction between the S2S3 loop of the channel voltage sensor and the EF3 of CaM (1, 21), which, in turn, is engaged to the AB fork. These structural studies suggest that the privileged role of EF3 may derive from constrains imposed by the channel architecture. To address the significance of this interaction in the absence of other channel domains, changes in the fluorescent emission of dansylated CaM (D-CaM) produced by peptides derived from the K_V_7 S2S3 sequence were monitored (38). Interaction of alpha helices within the groove of the CaM lobes results in an increase in fluorescent emission of D-CaM, whereas the binding of Ca^2+^ to the EF-hands causes, in addition to an increase in fluorescence, a leftward shift in the position of the peak in the emission spectrum (38).

The response to S2S3 peptides rendered an analogous profile to that of Ca^2+^: a leftward shift on the emission peak, and an increase in fluorescent emission (Fig 3). A similar responses were observed with S2S3 peptides derived from the sequence of human K_V_7.1 through K_V_7.5 (n = 3) (Supplemental Fig 5). The relative increase in emission intensity was twice as large in the absence of Ca^2+^ (Fig 3). This is in contrast to what has been observed for peptides or targets that are embraced within the CaM lobes, in which the relative increase is similar with and without Ca^2+^ (39, 40). Interestingly, the leftward shifts caused by Ca^2+^ and the peptide were additive (Fig 3). These results could be explained if Ca^2+^ and the peptide where interacting with CaM simultaneously. The displacement of the dose-response to the right (Fig 3B), increasing the EC_50_ value, suggest that Ca^2+^ mitigates the effect of S2S3 on D-CaM.

**Figure 5.**
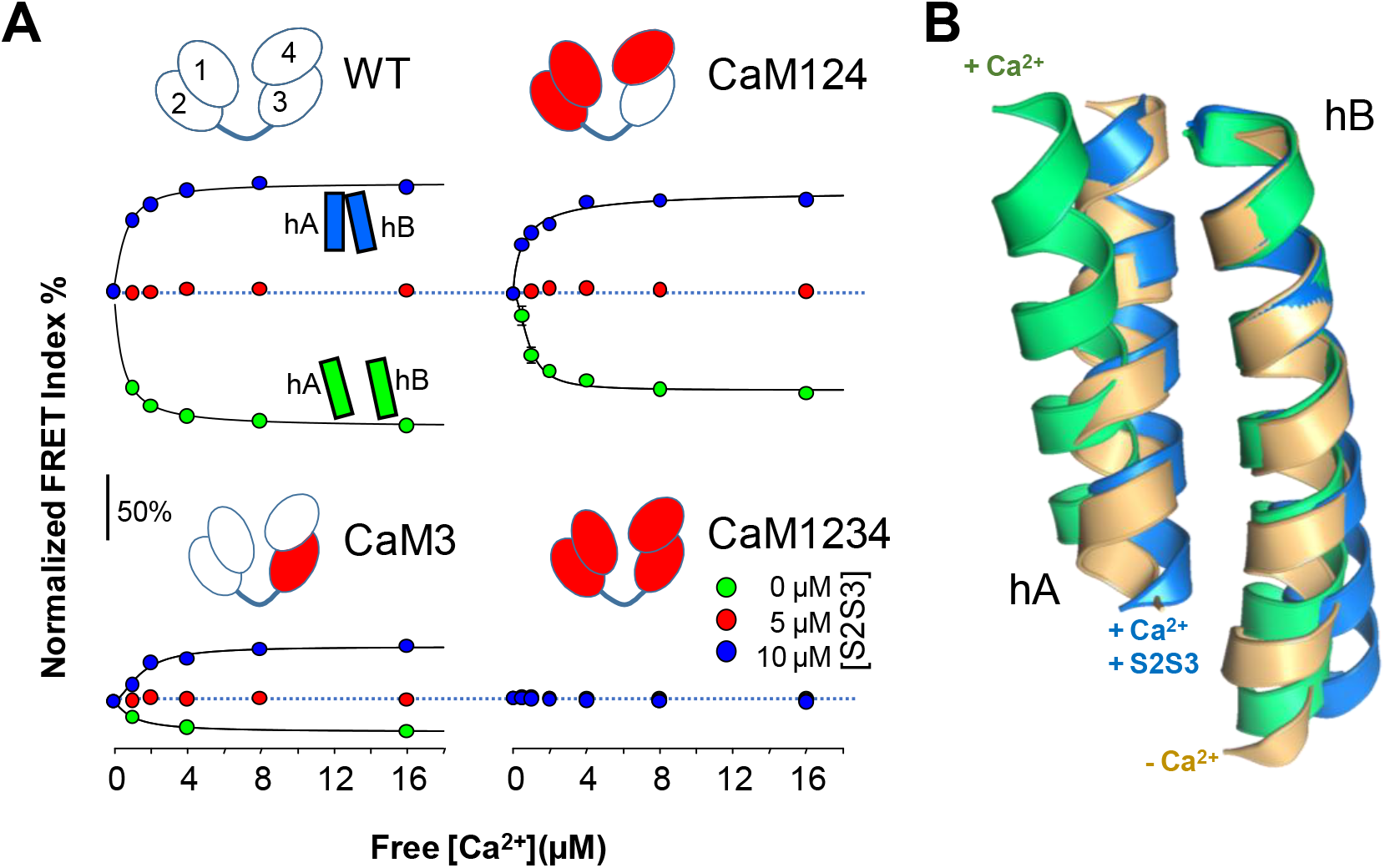
Relative FRET changes of the human K_V_7.2 in complex with mutant CaM. **A**. Ca^2+^ dose-response in the presence of the indicated concentrations of S2S3 peptide, normalized to the maximal response. FRET changes expanded from 100% reduction to 70% increase. Dotted lines mark no changes in FRET. The CaM mutant forming part of each complex is indicated on top of the corresponding dose-response curve. The continuous lines are the result of the fit of a Hill equation, with EC_50_ values of 0.84 ± 0.04 and 0.83 ± 0.07 µM in the absence of S2S3 for WT and CaM124 (n = 4), respectively. In the presence of 10 µM S2S3, the EC_50_ values were 0.62 ± 0.91, 0.62 ± 0.16 and 1.48 ± 0.22 µM for WT, CaM124 and CaM3, respectively (n = 4). **B**. Superposition of helices A and B solved in the absence of Ca^2+^ (gold, PDB 6FEH, K_V_7.2), in the presence of Ca^2+^ (green, PDB 6FEG, Kv7.2), and interacting with the S2S3 loop in the presence of Ca^2+^ (blue, PDB 5VMS, K_V_7.1).

### NMR reveals interaction of the S2S3 peptide with the C-lobe of the AB/CaM complex

The NMR signals from labeled WT CaM complexed with non-labeled K_V_7.2 AB fork were compared in the presence and absence of the S2S3 peptide, and with Ca^2+^ added (holo-CaM, four EF-hands Ca^2+^-loaded) or not added (int-CaM, N-lobe Ca^2+^-loaded). Chemical shift perturbations (CSP) produced by the S2S3 peptide in the ^1^H-^15^N-HSQC map of int-CaM (holo-N-lobe and apo-C-lobe) and holo-CaM in complex with the K_V_7.2 CRD are shown in Fig 4A (See also Supplemental Fig 6). In the presence of the S2S3 peptide, several resonances of CaM residues in the spectrum were shifted, most of them located in the C-lobe. The CSP perturbations, color-coded in the structure of the human K_V_7.2 CRD in Supplemental Fig 6, are consistent with the S2S3 loop interacting predominantly with the EF3 Ca^2+^ loop, both in absence and in the presence of Ca^2+^. EF3 displacements were observed for D94, N98, Y100, I102 and A104, whereas for EF4, changes in the environment of I131 and E139 are beyond the threshold level (Fig 4B). Thus, Ca^2+^ addition produces a significant perturbation map, which is in line with the differential relative increase in fluorescence caused by the peptide in the D-CaM assay (Fig 3B). Next, we performed atomistic molecular dynamics (MD) simulations to investigate the interactions between the K_V_7.1 S2S3 peptide and int- or holo-CaM in complex with the K_V_7.2 CRD. Consistent with the NMR interaction experiment, the contact map obtained from the simulations shows that the peptide interacts mainly with the EF3 loop and the linker connecting CaM lobes (Figs 4C-D and Supplemental Fig 7). In contrast, there were not contacts in the region connecting EF3 and EF4, suggesting that the CSPs observed are better interpreted as an allosteric effect, rather than a direct contact with the peptide.

**Figure 6.**
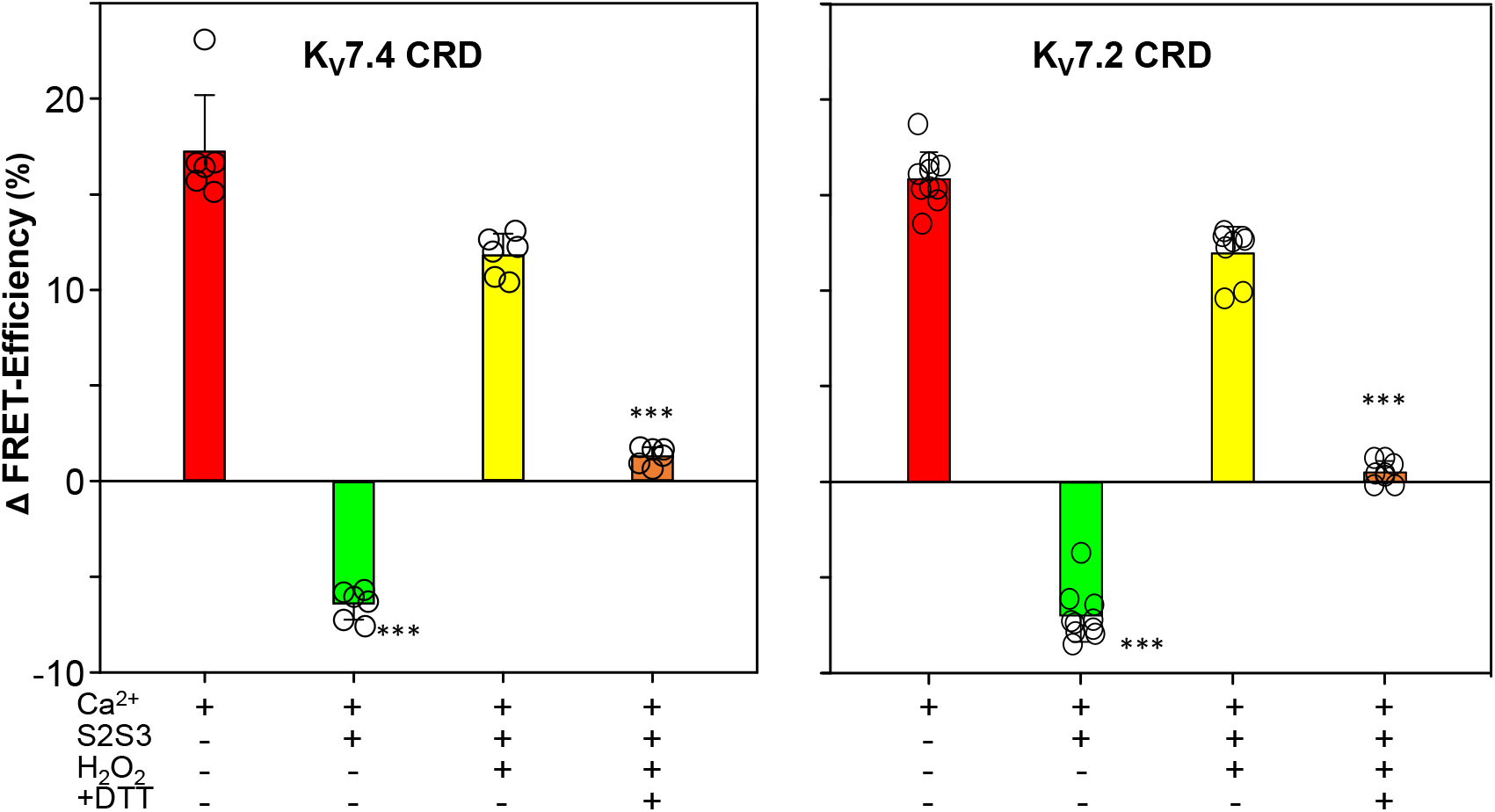
FRET efficiency changes prompted by oxidized and reduced S2S3 peptides. Difference in FRET efficiency in the absence and the presence of Ca^2+^ with sensors derived from the K_V_7.4 and K_V_7.2 CRD sequences. Red, sensor alone. Green, in the presence of 10 µM Q1-S2S3 peptide. Yellow, with 10 µM oxidized peptide. Orange, oxidized peptide treated with 1 mM DTT. Bars represent mean ± SEM FRET-efficiency. *p < 0.05; ***p < 0.001. Each plot represents the average of at least 6 independent experiments.

**Figure 7.**
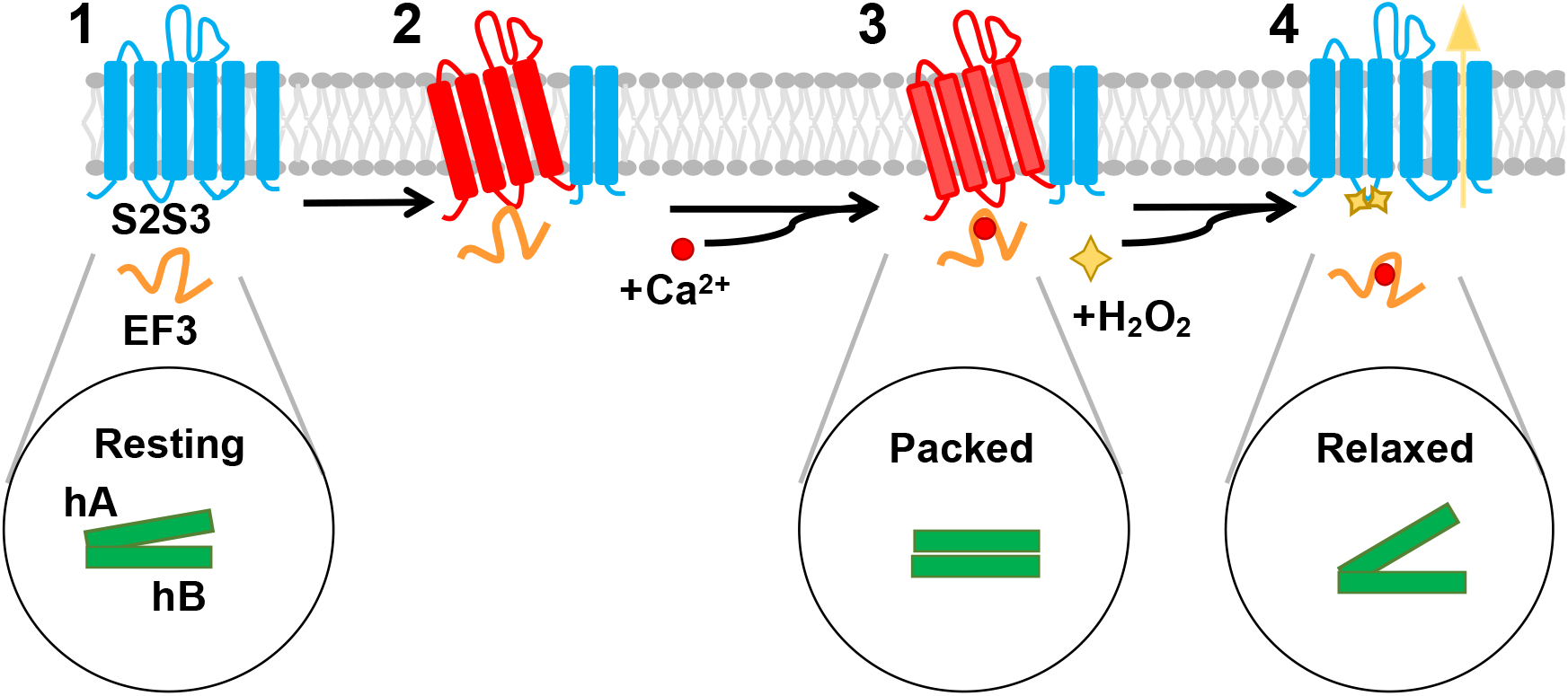
Model for K_V_7/CaM-mediated redox response. **[1]** Cartoon representation of the channel with the VSD engaged with the pore, unloaded EF3 away from S2S3, and the fork formed by helices A and B in a resting position. [**2**] EF3, by interacting with S2S3, stabilizes the state where VSD is disengaged from the pore. The S2S3 loop of the K_V_7 channel can interact with EF3 of CaM either in the absence [2] or in the presence of Ca^2+^ [3]. When EF3 interacts with the S2S3 loop in the presence of Ca^2+^, the CRD domain adopts a packed state, and channel opening is unlikely, resembling the “reluctant” state proposed for voltage-dependent Ca^2+^ channels (54). [**4**] The redox response of H_2_O_2_ is only manifested if Ca^2+^ is bound to EF3. H_2_O_2_ oxidizes the cysteine residues present in S2S3, favoring an engaged state of the channel, resembling the “willing” state proposed for voltage-dependent Ca^2+^ channels (54). In this state, a remarkable increase/recovery of M-current density in seconds is observed. The combination of changes on the relative orientation of hA and the gate, and the angle between hA and hB may facilitate or make more difficult the opening of the gate.

Regarding the S2S3 peptide, residues that form an intracellular loop located between W166-G176 are the ones that interact predominantly with CaM (Fig 4D). During the course of the simulation, the C-terminal region adopted an α-helix conformation for ≥ 97.8% of the time (see Supplemental Fig 8). The N-terminal that started as a 3_10_ helix became unstructured after the initial equilibration. It is reasonable to expect such differential stability since the N- and C-helices were initially formed by 6 and 10 residues, respectively, and 3_10_ helices are less stable than α-helices (41).

The interaction between S2S3 and EF3 was more stable when was not loaded with Ca^2+^ (Fig 4E, left). In contrast, the main contacts of the holo-system were established primarily with the linker connecting CaM lobes (Fig 4F, right). To analyze the interaction between S2S3 and EF3, we measured the distance between the center of mass of EF3 and S2S3 loops or the linker connecting the CaM lobes. The results suggest that the interaction between the S2S3 and the empty EF3 loops is rather stable, whereas Ca^2+^ occupancy prompts the movement of the peptide away from EF3 towards the linker on the lobes (Fig 4E). Thus, Ca^2+^ occupancy has an important influence on the S2S3/CaM interaction.

### Reversal of Ca^2+^-EF3 signaling by S2S3 peptides

Changes on FRET index in response to Ca^2+^ in the presence of S2S3 peptides were monitored as previously described (see Fig 2). Figure 5 shows that the Ca^2+^-dependent reduction in FRET index was mitigated as the concentration of peptide was increased. At high peptide concentrations (≥ 10 µM), the FRET index increased, suggesting that the distance/orientation of AB helices was even more favorable than in the absence of Ca^2+^. A similar behavior was observed when the effect of the peptide for the other K_V_7 family members was examined (Supplemental Fig 9).

The response to Ca^2+^ in the presence of S2S3, in terms of FRET index, was in the opposite direction than when the peptide was absent. The magnitude of signaling reversal was similar in WT and CaM124 complexes, whereas it was reduced in complexes with CaM3 (Fig 5). Thus, the direction/orientation of the movements in the AB fork when EF3 is loaded with Ca^2+^ is reversed upon interaction with S2S3.

### All K_V_7 CRDs display a similar response to Ca^2+^

A panel of K_V_7 biosensors in which the fork sequence was replaced by the equivalent segment from K_V_7.1 through K_V_7.5 human isoforms were created. FRET index was reduced for all biosensors in the presence of Ca^2+^, and the signal was significantly preserved in complexes formed with CaM124, whereas it was decreased in complexes with CaM3. The apparent affinity for Ca^2+^ was lower in the presence of the peptides, but the difference was not statistically significant (Supplemental Fig 9). Thus, we conclude that EF3 plays a similar role across the K_V_7 family of CRDs.

### Treatment with H_2_O_2_ reduces the effect of S2S3 peptides

The S2S3 loop, which is highly conserved among K_V_7 channels, contains one (K_V_7.1) or three cysteine residues (K_V_7.2-K_V_7.5). The cysteine site mediates an increase in open channel probability in response to oxidizing conditions (10). We tested the influence of oxidation by removing DTT from the buffer, and including H_2_O_2_ to obtain a derivate that will be referred to as oxidized-S2S3. There was a negligible impact of this treatment on helicity as assessed by circular dichroism (Supplemental Fig 6). Contrary to the increase observed with control S2S3 peptide, no changes in fluorescent emission of D-CaM were observed after addition of oxidized-S2S3 (Supplemental Fig 10).

The Ca^2+^ titration profile using K_V_7.2AB/CaM complex in the presence of oxidized S2S3 peptide (10 µM) was similar to that obtained in the absence of control S2S3, suggesting that oxidized S2S3 can no longer affect the AB-CaM interaction. FRET decreased from an efficiency of 40.7 ± 1.03 (n = 10) in the absence of Ca^2+^ to 30.5 ± 0.71 in the presence of 1 mM Ca^2+^ (n = 10). These values are similar to those obtained in the absence of control S2S3, where the efficiency of FRET dropped from 44.2 ± 0.03 (n = 6) to 29.1 ± 1.17 (n = 6). Using the K_V_7.4/CaM complex the behavior was almost identical. In the presence of oxidized peptide the FRET efficiency drops from 49.7 ± 0.67 (n = 16) to 41.4 ± 0.41 with Ca^2+^ (n = 6), which resembles the results obtained in the absence of control S2S3 peptide, where FRET efficiency decreases from 41.1 ± 0.40 (n = 6) to 27 ± 0.40 (n = 6) (Fig 5).

The oxidized S2S3 peptide was incubated with the reducing agent DTT aiming to reverse the effect of the treatment with H_2_O_2_. We observed a partial recovery: In the presence of reduced S2S3 peptide, FRET efficiency upon adding 1 mM Ca^2+^ reached a value of 43.3 ± 0.22 (n = 10) with the K_V_7.2 sensor and 43.3 ± 0.23 (n = 6) with the K_V_7.4 sensor (Fig 6). These results suggest that oxidation disrupts the interaction between S2S3 and EF3 loops.

## DISCUSSION

As its name suggests, calmodulin (CaM) is a CALcium MODULated protein, regarded as a fundamental player in the orchestration of Ca^2+^ signals in every eukaryotic cell (42). Notwithstanding, its function is not limited to Ca^2+^ signaling, having important functions in protein trafficking to the plasma membrane, protein folding, and other functions (43). Here, the portfolio of CaM capacities is extended by providing evidence of its essential role in transducing redox signaling in conjunction with K_V_7 channels, which exhibit an exquisite sensitivity to oxidation.

Previously, we demonstrated the significance of cysteine residues in neuronal K_V_7 channels located in the unusually long intracellular linker joining transmembrane segments S2 and S3, which are part of the voltage sensor (10). Here, we show that the ability of the EF3-hand of CaM to bind Ca^2+^ is essential in this redox signaling pathway. This is inferred, among others, from the observation that the effect of bath application of H_2_O_2_ is lost in cells overexpressing CaM variants with a disabled EF3-hand or treated with the membrane permeable Ca^2+^ chelator BAPTA-AM.

The redox response is characterized by a remarkable increase/recovery in M-current density on a second to minute time scale and it can be reversed by reducing agents (5, 10). This could derive from insertion of new channels, engaging silent channels, higher open probability, or a combination of these. Although the insertion/recruitment of new channels cannot be completely discarded, H_2_O_2_ causes an increase in single channel activity in excised patches were incorporation of new channels cannot take place (10). The redox impact on K_V_7 channels is accompanied by a left shift in the current-voltage relationship of macroscopic currents, meaning that channel opening becomes easier at lower voltages or that the probability of opening at a given voltage increases (10). Interestingly, K_V_7.3 channels, that present a very high open probability (44), do not respond to H_2_O_2_ (10), perhaps because there is little room for further channel activation.

Large leftwards shifts in voltage dependency of K_V_7 channels after over-expression of CaM C-lobe mutants (33, 45) have been reported, suggesting a critical role for EF3 (33). However, this shift has not always been observed (46). In this study, we saw relatively small (∼10 mV) but consistent leftward shift in voltage dependence of K_V_7.4 co-expressed with all CaM mutants containing the EF3 mutation, suggesting a degree of tonic channel inhibition conferred via EF3.

Taking into account the images at atomic resolution of CaM interacting with the voltage sensor of K_V_7.1 channels (1), it is tempting to propose that CaM makes it more difficult to reach the up position that leads to gate opening in response to depolarizations, or that CaM is stabilizing the state were the voltage sensor is disengaged from the pore. In other words, we envisage CaM constitutively and dynamically inhibiting channel activation and that the redox action relieves this inhibition by weakening the dynamic CaM-S2S3 interaction in a Ca^2+^-dependent manner. This idea fits with the observation that EF3 is not interacting with S2S3 in the available atomic resolution structures trapped in a partially (21) and fully (47) open states. To harmonize with other observations, we propose that this inhibition is counterbalanced by CaM-dependent promotion of surface expression when CaM or CaM1234 are over-expressed (48, 49). Further, our data suggest that Ca^2+^ binding to EF3 should help releasing the voltage sensor from CaM.

Interestingly, FRET changes in the isolated recombinant CRD caused by S2S3 peptides depend on the concentration of Ca^2+^, are reversibly sensitive to H_2_O_2_ treatment, and are primarily governed by EF3. The dose-response relationship with D-CaM and our MD simulations illustrate that the interaction is weaker when the EF3-hand is loaded with this cation. It is very clear that the influence of S2S3 on the relative orientation of helices A and B is only manifested in the presence of Ca^2+^, which is also a necessary condition for functional effects of hydrogen peroxides on K_V_7.4 currents. The lack of any impact on energy transfer on the CRD reveals that the interaction under low Ca^2+^ conditions does not result in a conformational change in the AB fork.

Remarkably, we find that EF3 is essential and sufficient within the isolated recombinant CRD to translate Ca^2+^ signaling into conformational changes, which result in an 18° opening of the AB fork (2). Thus, signaling through EF3 is a property inherent to the CRD, and does not derive from constrains that the geometry of the voltage sensor-pore imposes. We note that the architecture of K_V_7 channels allows exploiting EF3 signaling in a more efficient way. We call attention on the conditional duality of this signaling system. One branch operates on the voltage sensor (S2S3/EF3), and another branch changes the orientation of helix A (Fig 7), likely affecting S6, and therefore the main gate formed by the S6 bundle crossing.

An unexpected observation was the reversal in FRET index observed at higher S2S3 peptide concentrations only in the presence of Ca^2+^. Although our data suggest that S2S3 interacts with CaM in the presence and absence of Ca^2+^, occupation EF3 by this cation is a requirement to signal the reorganization of the CRD and for the functional redox effect on the channel. FRET changes caused by Ca^2+^ are opposed in the presence of S2S3 peptides. At the molecular level, we do not know what the reversal of the signal implies, because any modification in distance or orientation causes FRET alterations. It seems reasonable to propose that, in this scenario, the movement within the AB helices goes in the opposite direction, leading to a tighter packing of the fork, which is consistent with increased FRET signal. Tighter packing is what is observed when comparing the K_V_7.2 fork in absence of S2S3 -with and without Ca^2+^-with the S2S3 loop interacting with EF3, presumably loaded with Ca^2+^ (1) (Fig 5B). It has to be noted that various other modes of CaM induced modulation of K_V_7 channels were also proposed in the past, thus N-lobe of CaM was suggested to mediate effect of transient cytosolic Ca^2+^ elevation on K_V_7.4 (50) and complex CaM lobe switching with raising Ca^2+^ was also proposed (46). Hence, there is room for further research, especially with regards to the specific molecular interactions during rapidly changing cytosolic Ca^2+^ dynamics induced by Ca^2+^ signaling events in cells. Our data presented here, however, make a clear case for the specific importance of EF3 in bringing together redox- and Ca^2+^/CaM-dependent modulation of K_V_7 channel activity.

Based on docking calculations, the functional existence of significant interactions between a target protein and apo-CaM through the EF-hands of the C-lobe was first proposed for the smoothelin-like 1 protein (51). Subsequently, direct interactions between apo-EF3 and myosin were observed in X-ray structures, and this interaction was postulated to play an important role in signal transduction (52). Recent analysis of surface interaction from cryo-EM structures of ion channels with CaM hints to interactions between EF3 or EF4 with Eag1, TRPV5 and TRPV6 channels (53). Thus, Ca^2+^ signal bi-directional transduction through direct interaction between the Ca^2+^-binding sites and CaM target protein could be a widespread mechanism.

In summary, our data show that EF3 is essential for hA-hB FRET Ca^2+^ -dependent reduction and that Ca^2+^ and EF3 are essential for H_2_O_2_ effects on current potentiation (Figs 1, 2 and 5). We also show that S2S3 interacts with EF3, which alters Ca^2+^ effect on hA-hB FRET (Fig 5), Ca^2+^ reduces S2S3-EF3 interaction (Fig 4) and S2S3 oxidation by H_2_O_2_ eliminates the impact on Ca^2+^-dependent hA-hB FRET changes. From these data we infer that the S2S3-EF3 interaction is essential for the H_2_O_2_ effect, that oxidized S2S3 no longer interacts with EF3, and that interference with the S2S3-EF3 interaction led to current activation.

## MATERIAL AND METHODS

See SI material and methods.

## Acknowledgements

The Government of the Autonomous Community of the Basque Country (Elkartek BG2019, IT1165-19 and KK-2020/00110) and the Spanish Ministry of Science and Innovation (RTI2018-097839-B-100, RTI2018-101269-B-I00 and PID2021-128286NB-100) and FEDER funds. The work in NG’s laboratory was supported by Wellcome Trust Investigator Award (212302/Z/18/Z) and MRC Training grant (MR/P015727/1). E.N. and A.M-M. are supported by predoctoral contracts from the Basque Government administered by University of the Basque Country. C.M. was supported by the Basque Government through a Basque Excellence Research Centre (BERC) and J.U. was partially supported by BERC funds. C.D. thanks PRACE for awarding access to computational resources in CSCS, the Swiss National Supercomputing Service, in the 17^th^ and 20^th^ Project Access Calls. We acknowledge CESGA and CSIC for granting us access to computational resources to FinisTerrae II supercomputer.

## Notes

### Competing Interest Statement

The authors have declared no competing interest.

